# Sialophorin is an essential host element for vaccine immunity against pulmonary fungal infections

**DOI:** 10.1101/2022.03.31.486552

**Authors:** Srinivasu Mudalagiriyappa, George S Deepe, Som Gowda Nanjappa

**Author notes:** Corresponding author: Som G. Nanjappa.

## Abstract

The global burden of fungal infections is alarming, primarily due to the increasing immune-compromised population. The immuno-preventive/therapeutic measures, including vaccines, are necessary to prevent or control fungal diseases. Identifying a protective host element as a functional phenotypic marker is immensely valuable. We identified a host element, sialophorin, preferentially associated with antifungal memory T cells. We investigated its role in vaccine immunity using a mouse model of pulmonary fungal infection. We found that sialophorin was essential to bolster CD8^+^ T-cell responses to the vaccine by enhancing their differentiation and expanding cytokine-producing cells required for immunity. Using a gain-of-function approach, activating sialophorin using mAb augmented the CD8^+^ T cell responses, and sialophorin-sufficient CD8^+^ T cells were competitively superior in differentiation and expansion to the deficient cells. Sialophorin-mediated vaccine immunity was independent of the T cell trafficking effect. Finally, we show that sialophorin is a potential functional phenotypic marker of fungal vaccine-potency and immunity. Our study revealed that sialophorin is an essential host-target element to bolster vaccine responses and serves as a *potential biomarker* of fungal immunity.

**Author Summary:** Fungal infections have been rising in recent years due to increased immunocompromised individuals. Vaccination of *at-risk* individuals helps counter the infections. Thus, suitable vaccine platforms are needed with apt adjuvants, and a phenotypic marker of vaccine immunity will bolster the efforts. We identified a phenotypic marker, sialophorin, associated with T cell vaccine immunity to fungal infection. Our findings show an essential role of sialophorin for fungal immunity, as a target of adjuvanticity, and as a potential biomarker of vaccine immunity against many fungal infections.

## INTRODUCTION

Pathogen-host interactions in the host determine the course and outcome of the disease, including developing efficacious vaccine agents. Most fungal pathogens cause innocuous infections in a healthy individual but can be detrimental during immunosuppression. The increasing immunosuppressed population, including those with CD4^+^ T-cell deficiency, has elevated the incidence and severity of fungal infections in recent years [1–3]. In parallel, there is a surge in our understanding of the protective innate and adaptive host elements that spur the development of potent and efficacious fungal vaccine platforms [4]. Studies in recent years revealed the instrumental role innate immune cell elements such as CLRs and TLRs in shaping antifungal T cell responses [5, 6]. Similarly, cytokine receptors on T cells are either necessary or bolster the effector T cells to produce cytokines such as IL-17A, IFNγ, and GM-CSF, which activate innate immune cells including neutrophils and macrophages to kill pathogenic fungi [7–10]. Thus, the identity of a key T-cell element to enhance the vaccine responses, which can also be a potential biomarker of immunity against infections, can be a yardstick of antifungal immunity.

Sialophorin, first identified as a defective molecule in Wiskott-Aldrich Syndrome patients [11], is a glycoprotein expressed on T cells in two forms; low and high mol. wt., the latter a result of extensive O-glycosylation following cell activation. Existing literature suggest a negative role of sialophorin for primary T cell responses. Due to bulkiness and negative charges, sialophorin either has a modest effect on T cell responses or inhibits cell activation [12, 13]. Glycosylated sialophorin expression on activated T cells precipitates apoptosis, and thus the T cell memory pool is smaller. Low expression on T cells offers superior recall responses and immunity [14]. Sialophorin promotes T cell trafficking instigating encephalomyelitis [15]. Although reduced expression of sialophorin in Wiskott-Aldrich Syndrome patients with WASp mutation is associated with susceptibility to infections [16], the role of sialophorin for fungal immunity remains elusive. Here we elucidated the function of sialophorin in fungal vaccine immunity and its potential as a biomarker of fungal immunity.

## RESULTS

CD8^+^ T cells mount sterilizing vaccine-immunity to pulmonary fungal infection in CD4^+^ T-cell deficient hosts by expression of cytokines, including IL-17 (Tc17) lineage [17, 18]. Vaccine-induced antifungal CD8^+^ T cells develop as long-lasting memory without plasticity [19, 20]. We investigated a phenotypic marker associated with antifungal effector and memory CD8^+^ T cells and observed the upregulated expression of O-glycosylated form of sialophorin, CD43 (denoted gCD43 here onwards; Clone 1B11^+ve^; **Supplementary Fig. 1A**). The gCD43 is preferentially expressed on *polyclonal* Tc17 cells and a subset of *polyclonal* Tc1 (IFNγ ^+ve^) cells, but its expression may differ on the fungal antigen-specific T cells. To determine gCD43 expression on antigen-specific T cells, we developed an *in vitro* re-stimulation of antifungal CD8^+^ T cells with fungus-primed bone-marrow-derived dendritic cells (BMDC) as antigen-presenting cells. Most of the fungal-antigen responsive effector and memory IL-17^+^ CD8^+^ T (Tc17) cells expressed gCD43 (**Supplementary Fig. 1B**). Additionally, fungal antigen specific Tc1 cells showed an enriched gCD43 population. Thus, both effector and memory fungus-specific CD8^+^ T cells preferentially express gCD43.

Sialophorin (CD43) can be a marker of activation, such as CD44 [21], and may not have functional significance for fungal vaccine-immunity. To dissect this, we challenged vaccinated CD43^+/+^ and CD43^-/-^ mice to measure the fungal clearance in the lung. We found that CD43 was necessary for vaccine immunity to pulmonary fungal infection (**Fig. 1A**). Given that the extent of recall responses and antifungal immunity depends on quality and quantity of vaccine-induced responses, we measured the cytokine-producing CD8^+^ T cells at the peak of vaccine response.

**Figure 1.**
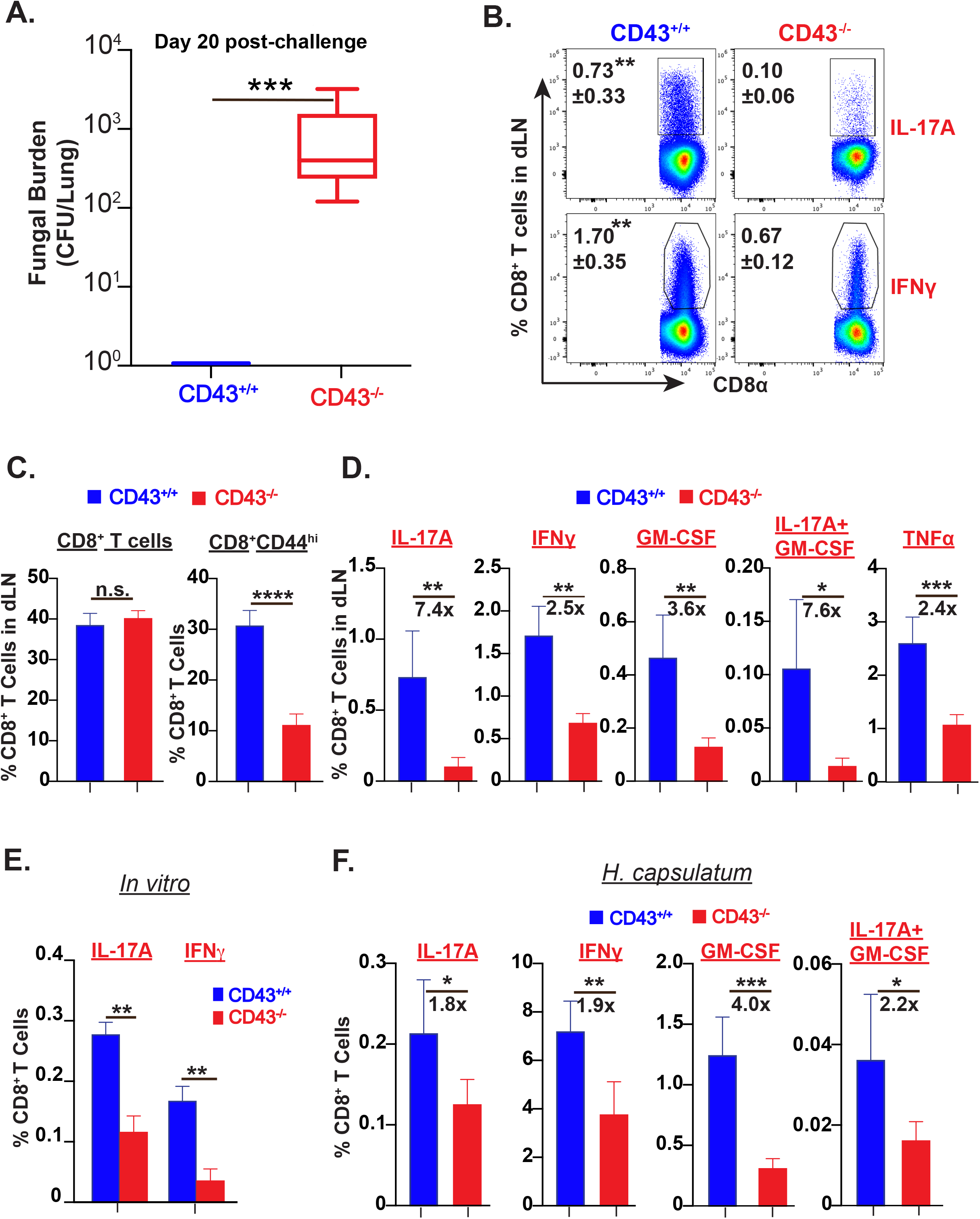
Sialophorin is necessary for fungal vaccine-immunity and induction of vaccine CD8^+^ T cell responses. (A) Sialophorin mediates vaccine-immunity. Naïve mice were vaccinated (Strain #55, ~1×10^5^ CFU, subcutaneously at dorsal and base of the tail), rested for six weeks, and intratracheally challenged with lethal strain #26199 (3×10^3^ CFU). On day 20 post-challenge, lungs were harvested to quantify fungal burden. CFU are in box and whisker plots. Data is representative of two independent experiments. N=6-7 mice/ group. p***≤0.001 vaccinated WT versus KO mice. (B-D) Sialophorin bolsters vaccine responses. The draining LN (dLN) and spleens were collected from vaccinated mice at ~3-wk post-vaccination. Single-cell suspensions were prepared for stimulation with anti-mouse CD3e (0.1μg/ml) and CD28 (1μg/ml) mAbs for 5hrs in the presence of GolgiStop (BD Bioscience). CD8^+^ T cells, their activation phenotype (CD44^hi^), and cytokine expressions were measured by flow cytometry. Data is representative of five independent experiments. Values are mean ± SD. N=4-5 mice/ group. p*≤0.05, p**≤0.01, p***≤0.001, and p****≤0.0001 vaccinated WT versus KO T cells. (E) Sialophorin enhances CD8^+^ T cell responses to fungal antigen *in vitro*. WT BMDC (1×10^6^ cells) were pulsed with heat-killed #55 yeast in a 6-well plate. A day later, enriched naïve CD8^+^ T cells were added (1 BMDC: 0.5 yeast: 2 T cells) and incubated for an additional 4 days. On day 5, GolgiStop (BD Biosciences) was added to the culture for 5 hours, and cells were washed, stained, and analyzed by flow cytometry. Data is representative of three independent experiments. Values are mean ± SD. N=4-6 replicates/group. p**≤0.01 WT versus KO CD8^+^ T cells. (F) Sialophorin augments vaccine responses to histoplasma. Naïve mice were vaccinated with *Histoplasma capsulatum* (10^6^ CFUs; subcutaneously) and at ~3-wk post-vaccination, single-cell suspensions from dLN were restimulated with anti-mouse CD3 and CD28 before analyzing cytokine^+ve^ T cells by flow cytometry. Data is representative of two independent experiments. Values are mean ± SD. N=5 mice/group. p*≤0.05, p**≤0.01, and p***≤0.001 T cells of WT versus KO mice.

We found severely impaired cytokine responses, strikingly of the Tc17 lineages, when CD43 was lacking, in cells from draining lymph nodes (dLN; **Fig. 1 B & D**) and spleen (**Supplementary Fig. 2**). Several considerations may explain the compromised responses in CD43-deficient cells. One possibility is that the immune response is biased to an unprotective Type 2 response induced by IL-4 or IL10 [22, 23]; however, neither were induced (data not shown). On the other hand, blunted responses in CD43-deficient mice may be caused by a reduction in CD8^+^ T-cell recruitment or developmental abnormality. The number of CD8^+^ T cells in control and CD43-deficient mice were similar (**Fig. 1C**). However, sialophorin potentiated the activation (**Fig. 1C & Supplementary Fig. 2B**) and differentiation (**Fig. 1D & Supplementary Fig. 2C-E**) of effector CD8^+^ T cells. To validate the *in vivo* results, we again exploited the *in vitro* system by priming BMDC to present fungal antigens to enriched naïve CD8^+^ T cells. We recapitulated the *in vivo* responses (**Fig. 1E**). To assess if it is true for another endemic fungus, we used a genetically related organism, *Histoplasma*, that induces predominantly Tc1 over Tc17 responses. CD43 was essential to augment the vaccine responses (**Fig. 1F**). Thus, CD43 is instrumental in the development of fungal vaccine responses and immunity.

The expansion phase of an adaptive immune response encompasses the activation, proliferation, and differentiation, often intertwined, following antigenic stimulation [24]. In a kinetic experiment following fungal vaccination, we asked if sialophorin governs T-cell activation, differentiation, or proliferation. The defective cytokine^+ve^ CD8^+^ T-cell profile was evident in the absence of CD43 at an earlier phase of a vaccine response (day 13 post-vac; **Fig. 2A**), and this defect become more pronounced later (**Fig. 2B**), suggesting an unceasing function of CD43 throughout the expansion phase of the vaccine response. In line with this, sialophorin enhanced the expression of lineage-defining transcription factors, RORγt and T-bet, which are essential to drive the differentiation process of T cells for type 17 and type 1, respectively (**Fig. 2D & Supplementary Fig. 3A**). CD43 promoted the proliferation of Tc17 to a greater extent than that of the Tc1 cell lineage (**Fig. 2G**).

**Figure 2.**
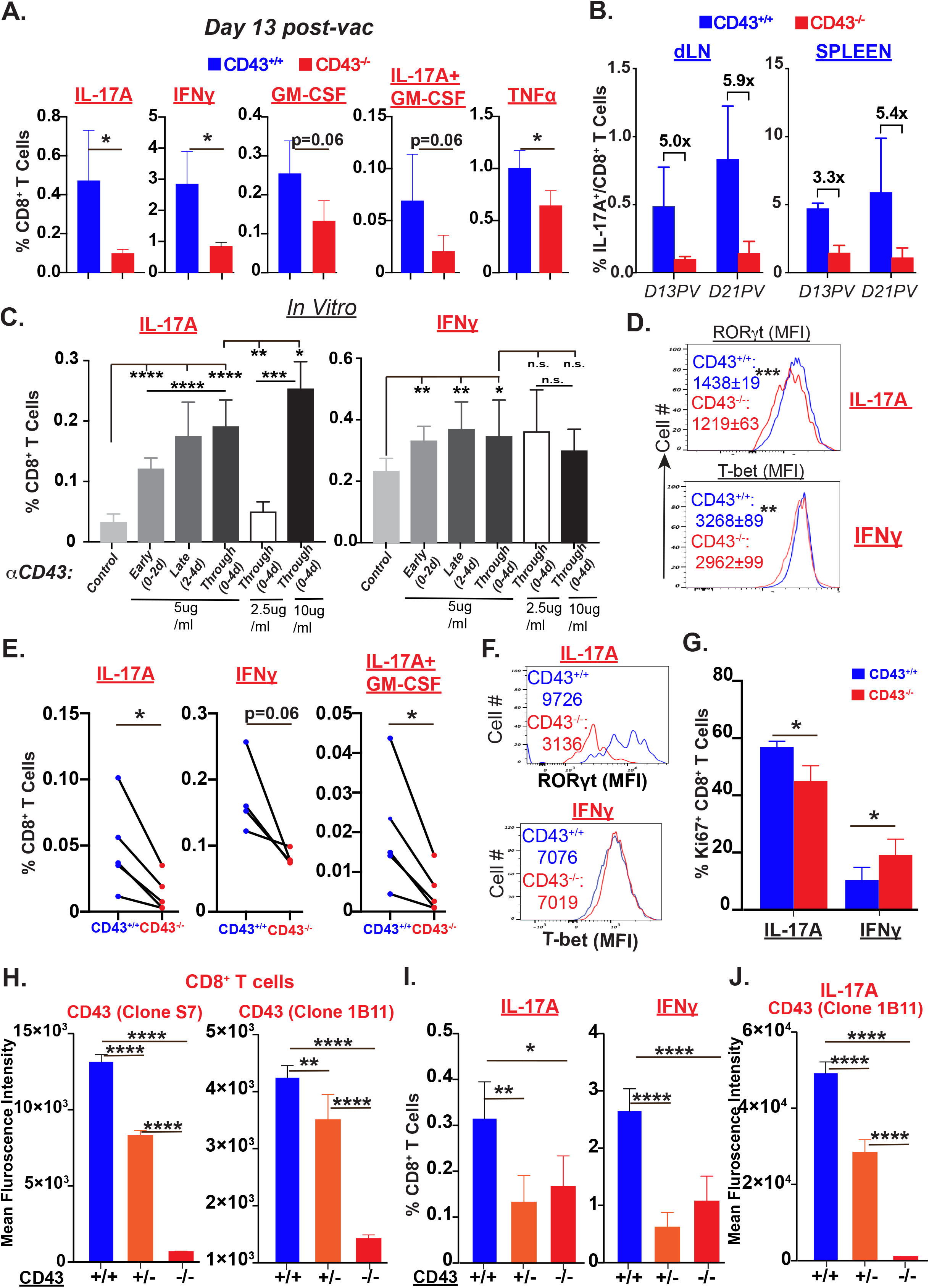
Sialophorin potentiates differentiation and expansion of CD8^+^ T cells. (A-B) Sialophorin is required throughout the expansion phase of T cell response. Cohorts of naïve mice were vaccinated, and at days 13 and 21, single-cell suspensions from dLN/spleen were restimulated with anti-mouse CD3e and CD28 mAbs for analysis of CD8^+^ T cell responses by flow cytometry by gating out dead cells. Data is representative of two independent experiments. N=4-5 mice/group. Values are mean ± SD. N=5 mice/group. p*≤0.05 of WT and KO T cells. (C) Stimulation of CD43 augments CD8^+^ T cell responses. Enriched CD8^+^ T cells from naïve CD43^+/+^ mice were incubated with yeast (heat-killed)-pulsed CD43^+/+^ BMDC and anti-CD43 (Clone 1B11) mAb was added for indicated days (N=4-6 replicates/group). CD8^+^ T cell responses were analyzed by flow cytometry. Data is representative of at least two independent experiments. Values are mean ± SD. p*≤0.05, p**≤0.01, p***≤0.001, and p****≤0.0001. (E) Cell-intrinsic role of sialophorin for CD8^+^ T cell responses to the vaccine. Enriched CD8^+^ T cells (1.4 x 10^6^) from naïve WT and KO mice were *co-transferred* (1:1) into TCRα^-/-^ recipients and vaccinated a day later. At ~3-wk post-vaccination, single-cell suspensions from dLN/spleen were restimulated with anti-mouse CD3e and CD28 mAbs for analysis of CD8^+^ T cell responses by flow cytometry. Data is representative of two independent experiments. Values are mean ± SD. N=5 recipients. p*≤0.05 of WT and KO T cells. (D, F, G) Sialophorin bolsters differentiation and proliferation of CD8^+^ T cells. Single-cell suspension cells, following restimulation, were surface and intracellular cytokine stained before staining for transcription factors/Ki67 staining using Transcription Factors Staining kit (eBioscience). Cells were analyzed by flow cytometry. The values of transcription factors are mean ± SD from (A) at day 13 post-vac and mean of pooled samples from ~3-wk post-vac (E). The percent values of Ki67 are mean ± SD from (E). N=4-5 mice/group. Data is representative of at least two independent experiments. p*≤0.05, p**≤0.01, and p***≤0.001 of WT and KO T cells. (H-J) Sialophorin expression determines the vaccine-induced CD8^+^ T cell responses. Naïve WT, Het, and KO mice were vaccinated, and at ~3-wk post-vaccination, single-cell suspensions from dLN/spleen were restimulated with anti-mouse CD3e and CD28 mAbs for 5hrs for analysis of CD8^+^ T cell responses by flow cytometry. Data is representative of two independent experiments. MFIs (H, J) and percent cytokine^+ve^ cells (I) values are mean ± SD. N=5 mice/group. p*≤0.05, p**≤0.01, and p****≤0.0001 of WT, Het, & KO T cells.

To validate these observations, we reasoned that use of an activating mAb against CD43 would enhance the CD8^+^ T cell responses. We, again exploited the *in vitro* fungal loaded-BMDC stimulation of the naïve T cells and used different concentrations of mAb (Clone 1B11) for different durations of stimulation, emulating an *in vivo* kinetic experiment. Exposure to the mAb enhanced both Tc17 and Tc1 responses in a dose and duration-dependent manner, exerting influence mainly on Tc17 cell lineage during late /throughout phases (days 2-4 or 0-4) of CD43 stimulation (**Fig. 2C**). Thus, a positive engagement of gCD43 is beneficial to augment antifungal T cell responses. Since BMDC also express CD43, we wanted to exclude the possibility that the mAb influenced T cell responses by activating BMDC. We deciphered the T cell-specific role of CD43 by adoptive *co*-transfers of naïve T cells before vaccination. This approach enabled us to test the specific role of CD43 for CD8^+^ T-cell responses, where the inflammatory milieu would be similar. In line with the earlier data, CD43-deficient CD8^+^ T cells exhibited lower cytokine responses than CD43-sufficient cells (**Fig. 2E**). Notably, the Tc17 lineage cells are more dependent on CD43 than the Tc1 cells. Similarly, the expression of lineage-defining transcription factor was significantly less in Tc17 cells than Tc1 cells under CD43 deficiency (**Fig. 2F**), suggesting the T-cell specific role of CD43 for their differentiation and expansion following fungal vaccination.

The *in vitro* mAb stimulation enhanced antigen-‘specific CD8^+^ T-cell responses (**Fig. 2C**), suggesting enhanced CD43 activation for bolstering vaccine potency; thus, we asked if partial CD43 expression will reduce vaccine efficacy. We used CD43 heterozygous mice to address this where one functional copy of the gene is present for examination. Heterozygous CD8^+^ T cells expressed reduced levels of both low (detected with mAb clone S7) and high mol. wt. gCD43 (**Fig. 2H**). Optimal vaccine responses of Tc17 and Tc1 cells required both copies of CD43. Cells with monoallelic CD43 phenocopied the blunted responses when both alleles were absent (**Fig. 2I**). The dampened responses by monoallelic cells were not caused by complete loss of CD43 on differentiated CD8^+^ T cells in heterozygous mice (**Fig. 2J**) or a decrease in the number of CD8^+^ T cells in the dLN (**Supplementary Fig. 3C**). Thus, both alleles of CD43 must be present for effective fungal CD8^+^ T cell responses.

T cell-dependent vaccine immunity requires the migration of effector or memory T cells to the site of infection to execute their functions against pathogens. While the threshold number of qualitatively superior effector T cells is necessary, the number of protective cytokine-producing T cells at the site of infection, in this case the lung, dictates the efficacy of vaccine immunity in pulmonary fungal infection. Following vaccination and pulmonary challenge, we measured the cytokine producing CD8^+^ T cells in the lung and assessed fungal burden at the peak of the anamnestic response (day 4-5 post-challenge). CD8^+^ T cells and their responses in the lung were vaccination-dependent (**Fig. 3C**), and fungal control was poor in unvaccinated controls (**Fig. 3A**). Vaccine immunity required sialophorin, which also reduced overt pathology associated with impaired fungal control (**Fig. 3A**). The deficiency of sialophorin dampened T-cell recall responses, especially of the Tc17 lineage, but not the Tc1 (**Fig. 3B-C & Supplementary Fig. 4A**) cells in the lung. Since the CD8^+^ T-cell recall responses required sialophorin (**Fig. 3B**) for vaccine immunity (**Fig. 3A**), and sialophorin was implicated in T cell recruitment, we asked if effector CD8^+^ T-cell trafficking into the lung depended on it following infection. To address this, we used the intravascular staining technique to differentiate vascular vs. parenchyma CD8^+^ T cells. We reasoned that if effector CD8^+^ T-cell trafficking requires sialophorin, there will be fewer lung parenchyma cells than lung vascular or spleen (harbor vaccine effector T cells) compartments. We found a similar *proportion* of Tc17 cells in different tissue compartments (**Fig. 3D**; left panels) and higher proportions of Tc1 cells in the lung’s parenchyma and vascular compartments but not the spleen (**Fig. 3D**; right panels) in sialophorin-deficient mice. Thus, a minimal role of sialophorin for effector CD8^+^ T-cell trafficking into the lung. However, the trafficking of CD8^+^ T cells or their activated phenotype bearing CD44^hi^ cells into the lung was largely intact and did not depend on sialophorin (**Fig. 3E**).

**Figure 3.**
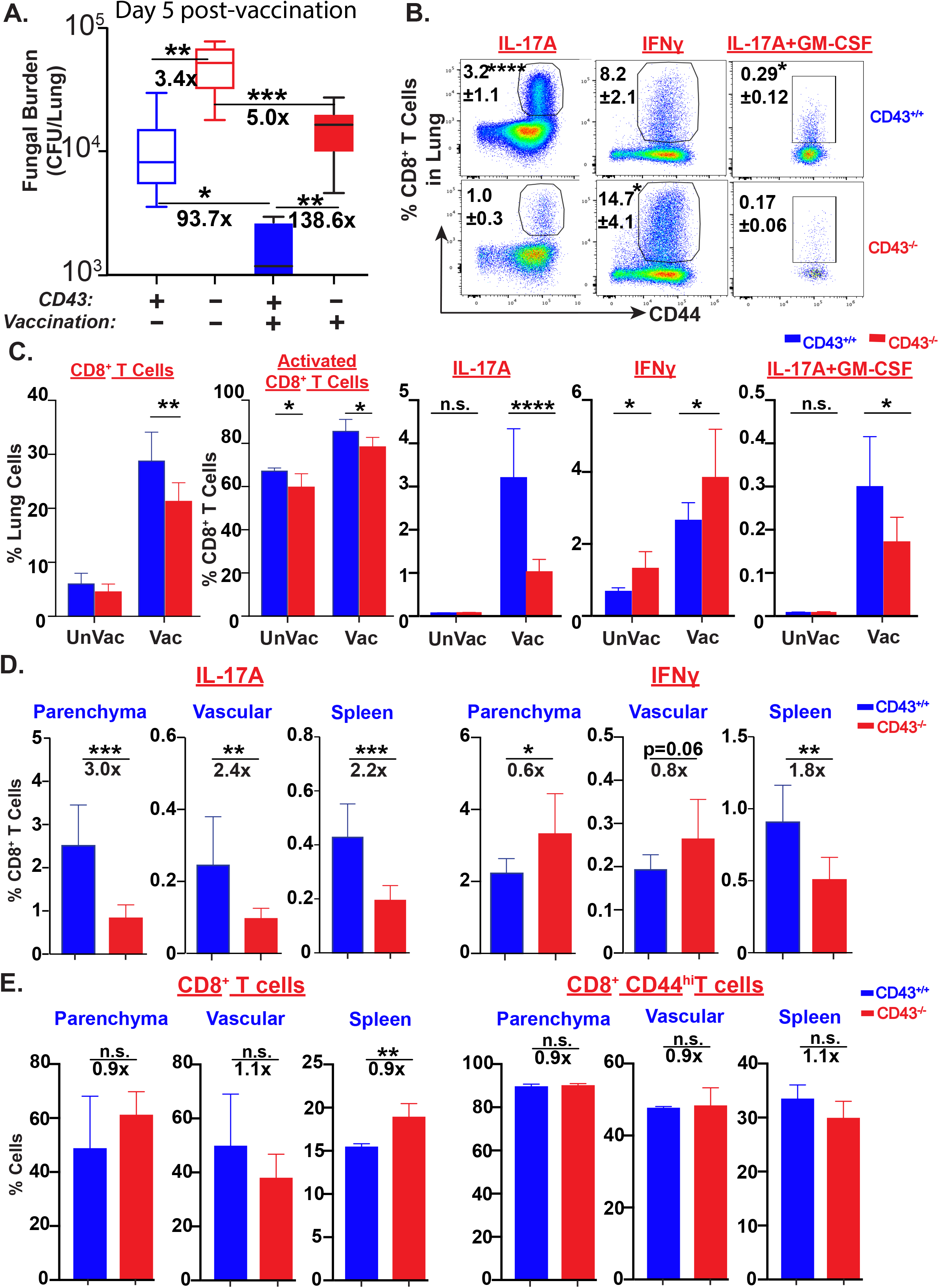
Sialophorin is necessary for CD8^+^ T-cell recall responses and immunity independent of its role in trafficking. (A) Sialophorin is required for vaccine-mediated immunity. Naïve CD43^+/+^ and CD43^-/-^ mice were vaccinated, rested for ~6 weeks, and intratracheally challenged with lethal yeast #26199 along with unvaccinated controls. On day 5 post-challenge, lungs were collected, homogenized, and plated on BHI agar plates to enumerate CFU. Data is representative of two independent experiments. N=5-8 mice/group. p*≤0.05, p**≤0.01, and p***≤0.001 of unvaccinated and vaccinated WT/KO mice. (B-C) Sialophorin is required for CD8^+^ T-cell recall responses. Single-cell suspensions from lungs of cohorts of vaccinated and challenged mice (A) were restimulated with anti-mouse CD3e and CD28 mAbs for 5hr and CD8^+^ T cell responses were analyzed by flow cytometry. Data is representative of two independent experiments. Values are mean ± SD. N=5 mice/group. p*≤0.05, p**≤0.01, and p****≤0.0001 of T cells of unvaccinated and vaccinated WT/KO mice. (D-E) Sialophorin is dispensable for effector CD8^+^ T-cell trafficking into the lung. Cohorts of vaccinated and challenged mice (A) were administered with fluorochrome-conjugated CD45.2 intravenously 3 min (intravascular staining) before the sacrifice. Single-cell suspensions from the lungs and spleen were restimulated with anti-mouse CD3e and CD28 mAbs for 5hr. Cells were stained for surface markers along with disparate fluorochrome-conjugated CD45.2 before staining for intracellular cytokines. CD8^+^ T cells and responses were analyzed by flow cytometry. Data is representative of two independent experiments. Values are mean ± SD. N=5 mice/group. p*≤0.05, p**≤0.01, and p***≤0.001 of WT and KO T cells.

Specific T-cell-dependent vaccine immunity against pulmonary fungal infection involves cytokine responses, which bolster innate cells activation to kill the yeast efficiently [8, 25]. Our data thus far suggested CD43, a transmembrane protein, is a functional phenotypic marker of fungal vaccine immunity. To elucidate this, we conducted classic vaccination strategies. We immunized mice with yeast-cell extract (YCE) antigens and heat-killed yeast adjuvanted with alum. After a boost and pulmonary challenge, we examined fungal control in mice and tested if it directly correlates with the CD43^+^ CD8^+^ T cells in the lung. Because vaccinated mice control fungal infection and sialophorin was necessary (**Fig. 3**), we hypothesized that effector Tc17 cells, which are essential for vaccine immunity [18], during recall responses preferentially express gCD43 and correlate with the fungal control. While, alum-adjuvanted YCE immunization provided significant immunity compared to unvaccinated controls, heat-killed yeast immunization delivered superior immunity (**Fig. 4 A**). The effectiveness of vaccination directly correlated with the proportion of CD43^+^ CD8^+^ T cells (**Fig. 4B**) and their IL-17A expression (**Fig. 4C**). Vaccination with heat-killed yeast alone provided a better response and immunity than the alum adjuvanted preparation. Immunity, including that induced by vaccination, against pulmonary fungal infection includes neutrophil activation to kill yeast [18, 26, 27]. The activation status, as assessed by CD11a, but not the numbers, of neutrophils positively correlated with the degree of Tc17 responses and CD43^+^ CD8^+^ T cells (**Fig. 4D**).

**Figure 4.**
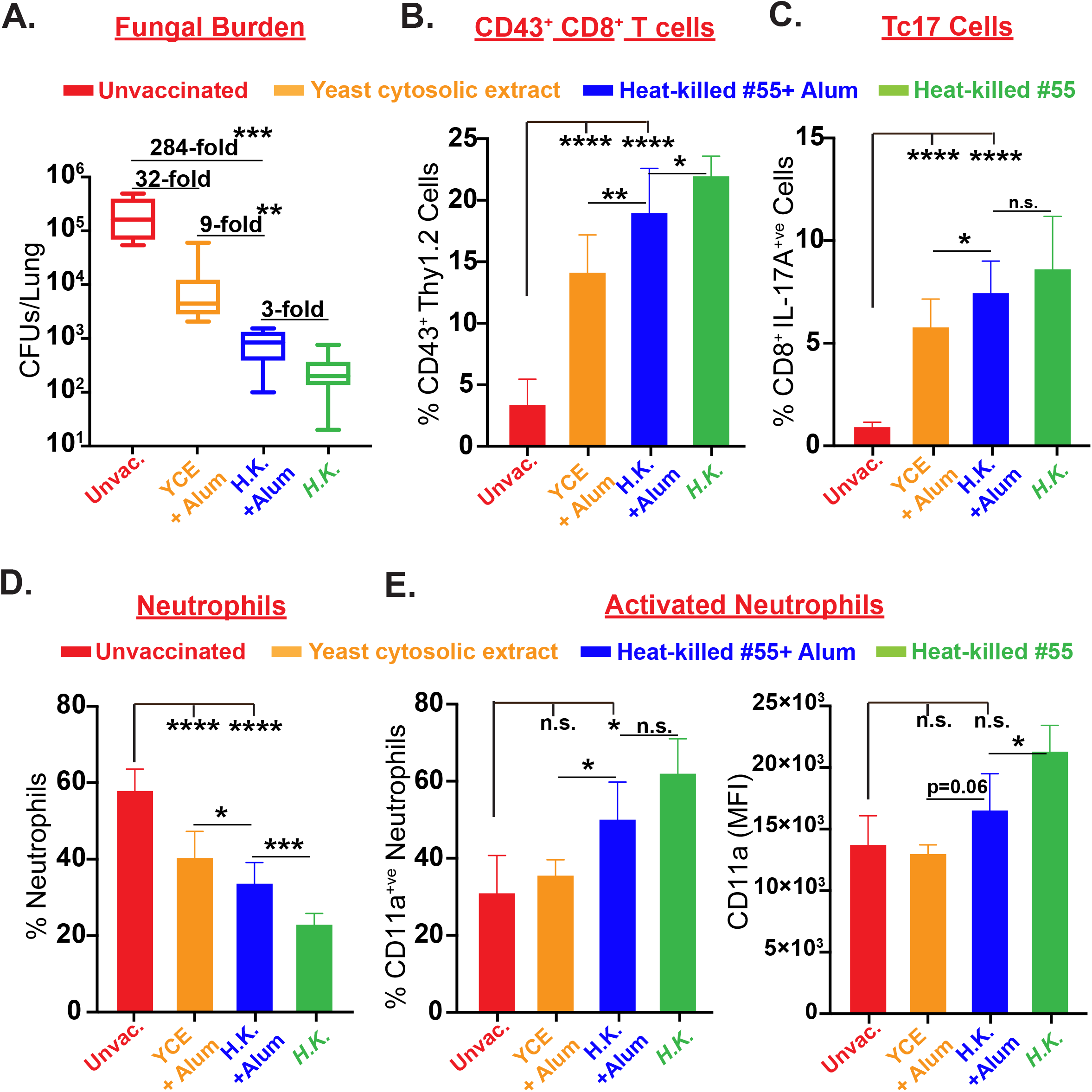
Sialophorin can serve as a functional phenotypic marker for anti-fungal immunity. (A) Fungal antigen vaccination elicits protective immunity. Cohorts of naïve CD43^+/+^ mice were vaccinated with ~150μg protein equivalent of yeast cell antigen or heat-killed yeast #55 with or without adjuvant, Alum. Mice were given booster dose once after two weeks and after a rest of 2 more weeks, animals were challenged intratracheally with virulent strain #26199 (5×10^3^ CFU). On day 5 post-challenge, lungs were harvested, homogenized, and plated on BHI agar to quantify fungal load. Data is representative of 2 independent experiments. N=5-10 mice/group. (B-C) Vaccine immunity is directly correlated with the recruitment of CD43^+^ T cells in the lung. Lung single-cell suspensions from infected mice (A) were restimulated with anti-mouse CD3e and CD28 mAbs for 5hr in the presence of GolgiStop (BD Bioscience). CD43^+^ and cytokine^+ve^ CD8^+^ T cells were analyzed by flow cytometry. Data is representative of 2 independent experiments. Values are mean ± SD. N=9-10 mice/group. p*≤0.05, p**≤0.01, and p****≤0.0001 of T cells among unvaccinated versus vaccinated groups. (D-E) Vaccine immunity is associated with the activation of neutrophils. Lung single-cell suspensions from infected mice (B-C) were directly surface stained for neutrophils and their activation phenotype (Live^+^, Thy1.2^−^, CD11b^+^, Ly6G^+^, CD11a^hi^). Data is representative of 2 independent experiments. Percent and MFI values are mean ± SD. N=9-10 mice/group. p*≤0.05, p***≤0.001, and p****≤0.0001 of neutrophils among unvaccinated versus vaccinated groups.

## DISCUSSION

Here we describe sialophorin as a host functional phenotypic marker for fungal immunity. Sialophorin deficiency blunted antifungal CD8^+^ T cell responses and engaging it with agonistic mAb enhanced the responses. Vaccine responses of both Tc1 and Tc17 subsets required sialophorin, which enhanced the expression of lineage-defining transcription factors and Tc17 cell proliferation. Unlike viral infections [13], our study showed that fungal-specific CD8^+^ T cells preferentially retained the O-glycosylated sialophorin expression during the memory phase and correlated with the immunity.

Despite the evidence that sialophorin functions as a negative T cell regulator, it can promote TCR-dependent Lck activation [28], enhance cell proliferation [29], increase cell survival [30, 31], and decrease the T-cell activation threshold [32]. In contrast, it is associated with a reduced memory T cell pool and poor recall responses to control viral or bacterial infections [14, 33]. We show here that sialophorin is essential for potentiating fungal-vaccine T cell responses, acting as a costimulatory molecule, and becoming a predominant phenotypic marker during memory homeostasis. Additionally, our data, in a vaccine platform, showed that glycosylated sialophorin expression on T cells directly correlated with expression of antifungal cytokines, activation of neutrophils, and fungal immunity.

T-cell trafficking to non-lymphoid organs, including the infection site, is a unique, essential feature of effector or memory T cells for immunity, and is mediated through adhesion molecules [34, 35]. Sialophorin facilitates Th1 and Th17 [36, 37] trafficking into tissues by interacting with E-selectin, a receptor for other ligands. Our data suggest the dispensability of sialophorin for Tc17, including Tc1, cell-trafficking into the lung following pulmonary challenge. We postulate the compensatory role of other ligands, including CD44 (**Supplementary Fig. 4B**), for Tc17 cell trafficking into the lung.

In line with our efforts to combat fungal infections, an increase in the knowledge of the mechanisms of immunity paved for several potential vaccine candidates [4]. Identity of a functional phenotypic marker that portends the potency of the fungal vaccine help in the success of the vaccination platform. Our study identified a surface molecule, sialophorin, essential for the vaccine responses and as a marker of immunity, especially of type 17 mediated.

## Materials and Methods

### Mice

C57BL/6 mice were obtained from Jackson Laboratories (Bar Harbor, ME) or Charles River (Wilmington, MA). Breeding pairs of B6.129S2-Tcra^tm1Mom^/J (TCRα KO) and B6.SJL-*Ptprc*^*a*^*Pepc*^*b*^/BoyJ (CD45.1) mice were purchased from Jackson Laboratories. Congenic CD45.1 mice were backcrossed with CD43^-/-^ mice (kindly provided by Pilar Alcaide, Tufts University) to generate congenic littermate controls. Mice were bred and housed in a specific-pathogen-free environment. All the experiments were performed using 6- to 8-week-old male and female animals. Animal housing and experiments were done according to strict guidelines of the Institutional Animal Care and Use Committee at the University of Illinois Urbana-Champaign (IACUC # 20221).

### Fungal Culture

Wild-type virulent *Blastomyces dermatitidis (ATCC strain #26199)* was purchased from ATCC. The isogenic strain of *Blastomyces dermatitidis* lacking BAD1 (Strain *#*55) was kindly provided by Bruce Klein (University of Wisconsin-Madison). Both the strains were grown on Middlebrook 7H10 agar slants supplemented with oleic acid-albumin complex (Sigma-Aldrich) maintained at 39°C in a humidified incubator. *Histoplasma capsulatum* (strain G217B) was grown in Histoplasma Macrophage Medium (HMM) for 72h at 37°C to vaccinate mice (~1 x 10^6^ CFUs).

### Vaccination, infection, and quantifying fungal burden

The mice were vaccinated subcutaneously (~1×10^5^ CFU of strain #55 or 100μg of fungal antigen prep) at two sites, the dorsal and the base of the tail regions. Fungal antigen prep-inoculated mice were given a booster dose after two weeks. For pulmonary infection or challenge, a virulent strain of Blastomyces *dermatitidis* (#26199, ATCC) was used (3-5×10^3^ CFUs) for intratracheal inoculation. Homogenized lung tissue samples were plated on brain heart infusion (BHI; Difco) agar to enumerate the fungal burden (CFUs).

CD4^+^ T cell depletion A weekly dose of 100 μg of mAb GK1.5 (BioXCell, West Lebanon, NH) by intravenous route was used to deplete CD4^+^ T cells with an efficiency of >95% depletion [19].

### Antibodies and flow cytometry

All antibodies were purchased from BD Biosciences, BioLegend, and Invitrogen. The single-cell suspensions were prepared from the tissues harvested on indicated days, and red blood cells were lysed using RBC lysis buffer. Draining lymph nodes (dLN) and spleen tissues were harvested to measure vaccine responses. Following pulmonary infection, the lungs were harvested to measure recall responses. Disparate fluorochrome-labeled anti-CD45.2 mAb was used for vascular staining (intravenous, 2ug/mouse, 3 mins prior to sacrifice) to distinctly mark circulating vs. parenchymal cells. Cells were restimulated with anti-mouse CD3e (clone 145.2C11; 0.1 μg/ml) and CD28 (clone 37.51; 1 μg/ml) antibodies in the presence of Golgi-stop or Golgi Plug (BD Biosciences) at 37°C for 5 hrs. Cells were incubated with anti-CD16/32 (Fc Block) antibodies and Fixable Live/Dead stain (Invitrogen) for 15 min before staining with antibodies for surface markers in 2% BSA/PBS buffer for 30 mins on ice. Following fixation using Fixation/Permeabilization kit (BD Biosciences) for 20 mins, cells were washed and stained with intracellular cytokine antibodies in 1X Perm/Wash buffer (BD Biosciences) for 45 mins at 4°C. For nuclear factors staining, cells were further fixed with Transcription Factor Fixation/ Permeabilization solution (eBioscience) before incubating with transcription factor/Ki67 antibodies. Cells were analyzed by flow cytometry using Cytek Aurora after excluding dead cells and data was analyzed with FlowJo v10.8.1.

### Adoptive transfer experiments

CD8^+^ T cells were enriched from naïve donor mice (CD43^+/+^ and CD43^-/-^) using BD IMag^TM^ Mouse CD8 T Lymphocyte Enrichment Kit (BD Bioscience) and equal numbers of cells were transferred into TCRα^-/-^ recipients.

*In vitro* stimulation of CD8^+^ T cells using bone marrow-derived dendritic cells Bone marrow (BM) cells were harvested from femurs and tibia of 6-8-wk old and cultured in RPMI media supplemented with 10% fetal bovine serum, strepto-penicillin, 20 ng/ml of GM-CSF, 10 ng/ml of IL-4, and 50 mM of mercaptoethanol at 37°C for six days. On day 7, BM Dendritic Cells (BMDC) were collected and incubated with heat killed *B.d.* #55 yeast for a day before co-incubation with enriched naïve CD8^+^ T cells (1 DC:0.5 yeast: 2 T cells) for additional four days. For restimulation to evaluate antigen-specific CD8^+^ T cell responses, yeast-pulsed BMDC were co-incubated with enriched effector or memory CD8^+^ T cells overnight. Golgi-stop (BD Bioscience) was added at the last 5 hours of stimulation, and cells were collected to analyze cytokine-producing CD8^+^ T cells by flow cytometry.

### Yeast antigen preparation

Yeast cytosolic proteins of *Blastomyces dermatitidis* (#55) were extracted as previously described [38, 39]. Strain #55 was grown on slants and harvested in phosphate-buffered saline (PBS; pH7.2) and incubated with thimerosal (1:10000) for 1 hr at room temperature. After washing, pelleted yeast cells were suspended with PBS containing inhibitors (0.5M EDTA,5μM Leupeptin, 200mM of PMSF) with 1:2 volume and subjected to bead beater for 6 mins with alternating 30s of pulsing and cooling at 4° C. Disrupted cells were centrifuged at 11,000g for 5mins, and the collected supernatant was yeast cytosolic extract (YCE) that was dialyzed with PBS for 36hr at 4°C before freezing at −70°C for further use.

### Statistical analysis

Statistical significance of differences in fungal lung CFU was measured by non-parametric Kruskal-Wallis (one-way ANOVA). A two-tailed unpaired Student’s t-test was used to determine statistical significance using GraphPad Prism 9.2 (GraphPad Software, LLC). A two-tailed P value of ≤0.05 was considered statistically significant.

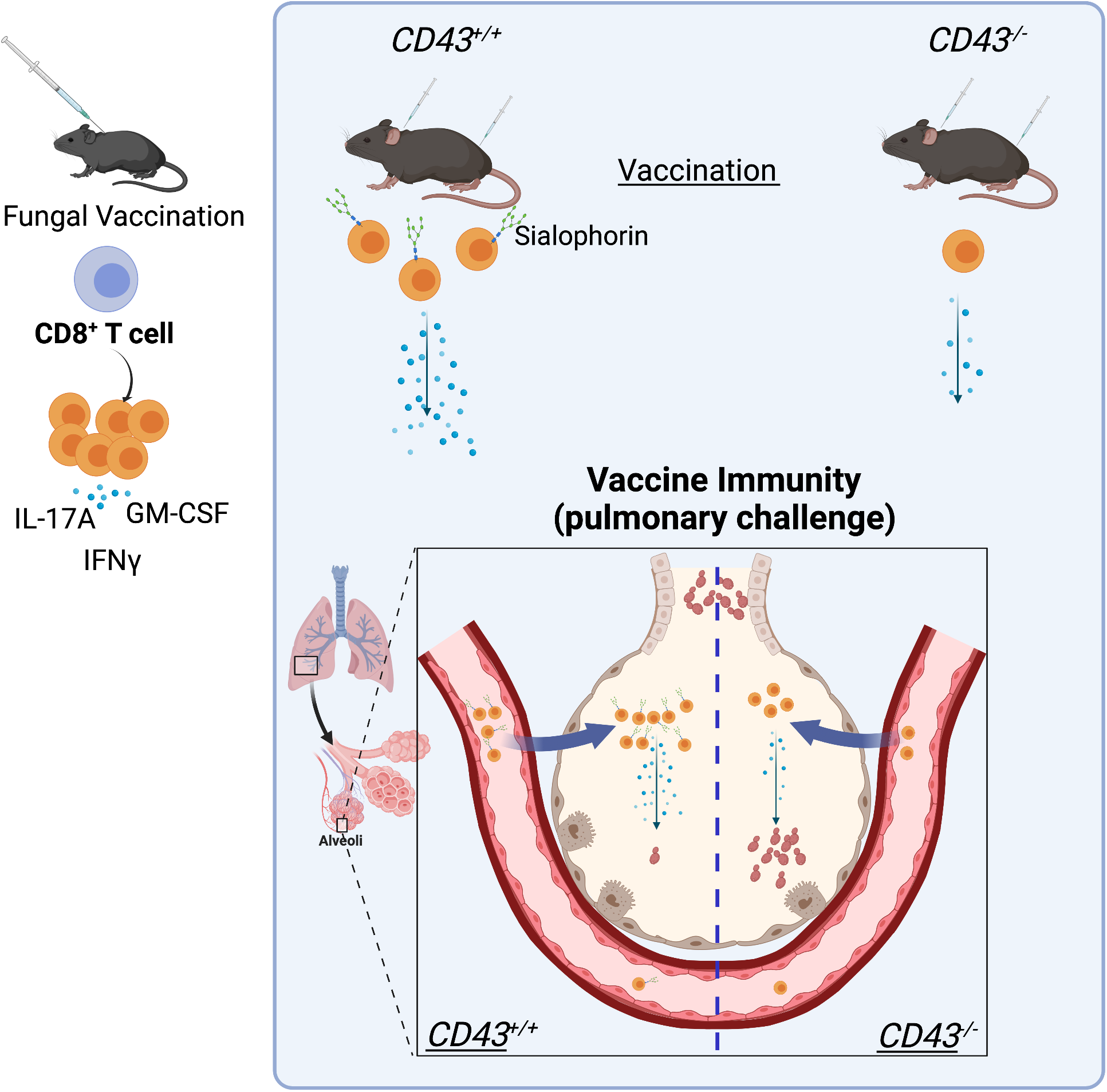

## Supporting information

Support Figure 1

Support Figure 2

Support Figure 3

Support Figure 4

## Acknowledgments

We sincerely thank Pilar Alcaide, Tufts University, for providing CD43^-/-^ mice and Bruce Klein, University of Wisconsin-Madison for providing *Blastomyces dermatitidis* #55 strain. We thank flow cytometry facility at College of Veterinary Medicine at the University of Illinois Urbana-Champaign. This study is supported by NIAID-NIH R01 AI153522 (SGN).

## Author contributions

SGN conceived the project. SGN & SM designed, executed, and analyzed the experiments. GSD provided reagents, designed, and helped analysis of histoplasma experiments. SGN and SM wrote the manuscript. SGN, GSD, and SM edited the manuscript.

