## Supplementary figures and images for "Sialophorin is an essential host element for vaccine immunity against pulmonary fungal infections"

### Support Figure 1

**Supplementary Figure 1**

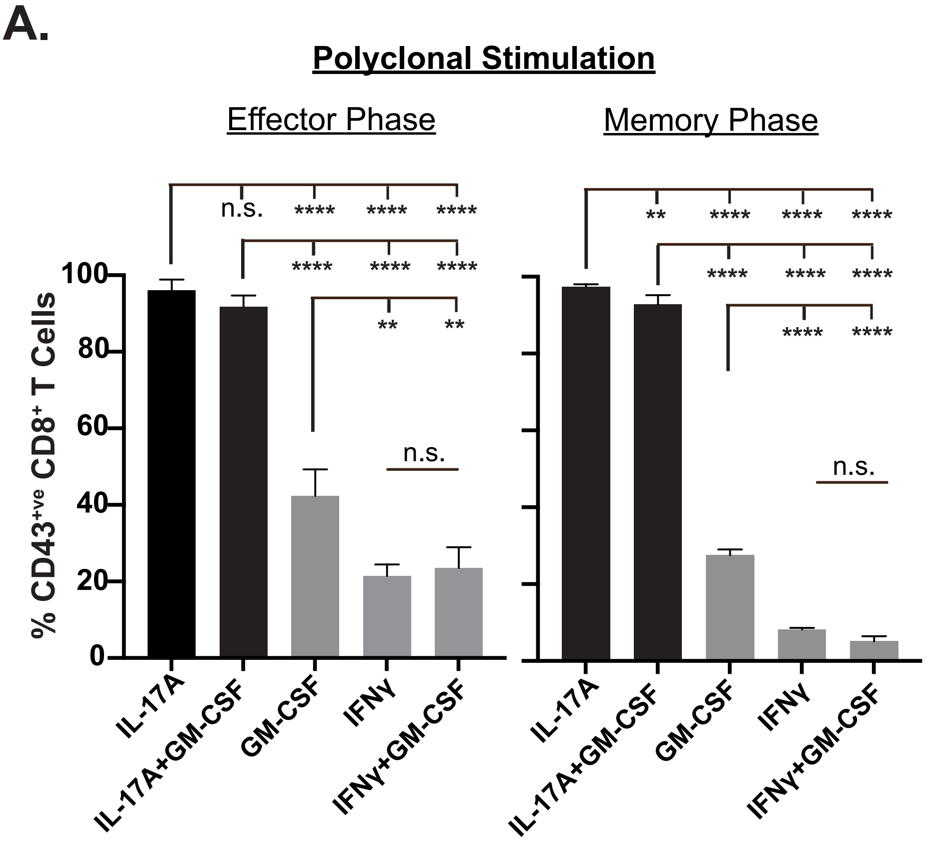


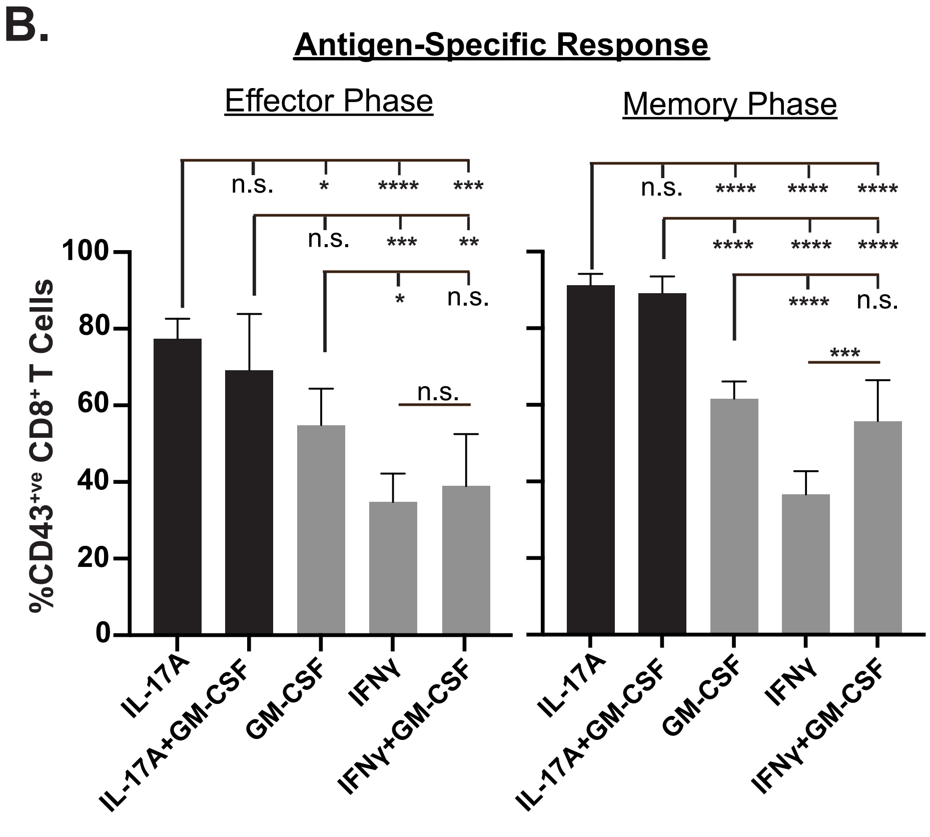

### Support Figure 2

Supplmentary Figure 2
related to Figure 1.

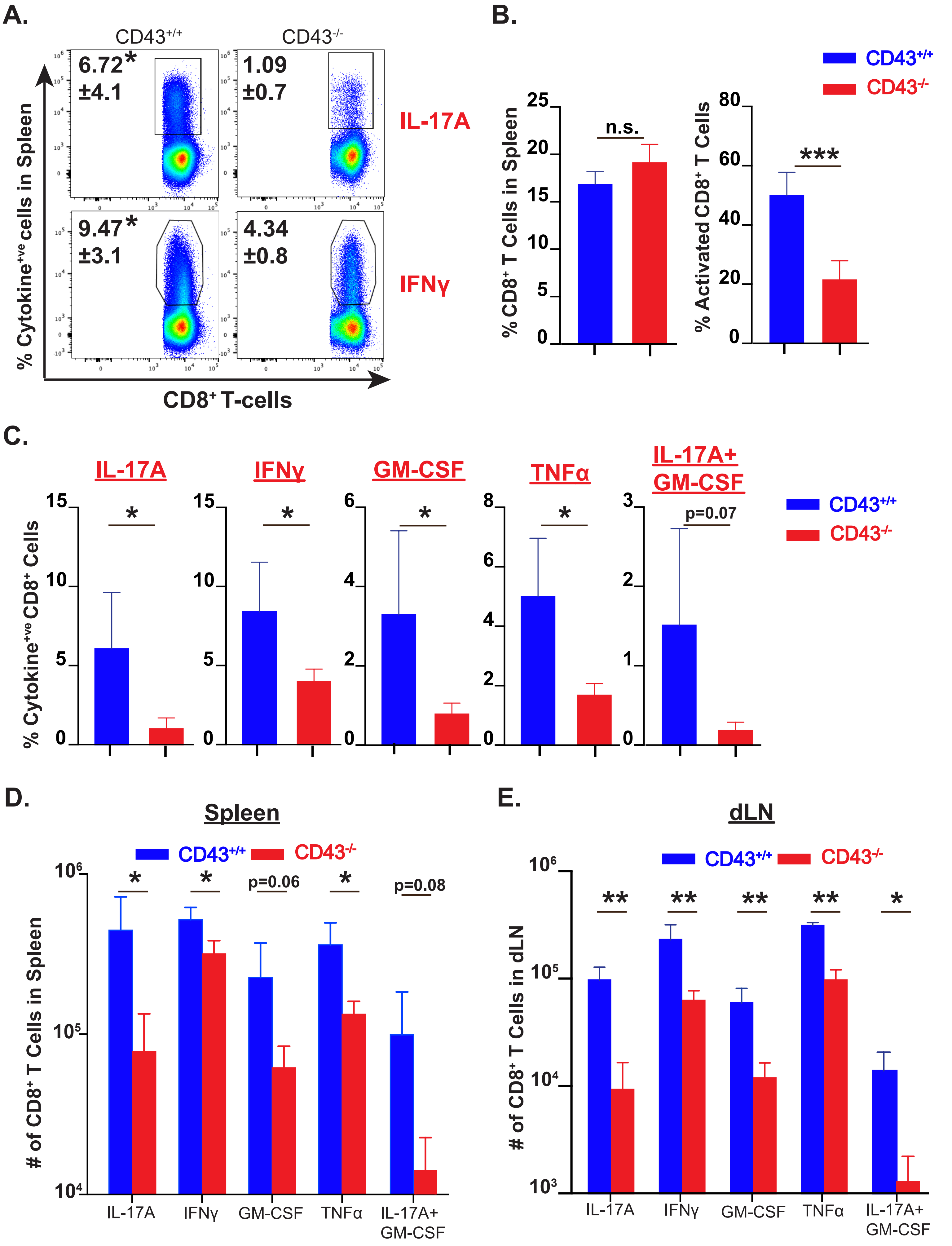

### Support Figure 3

#### Supplementary Figure 3

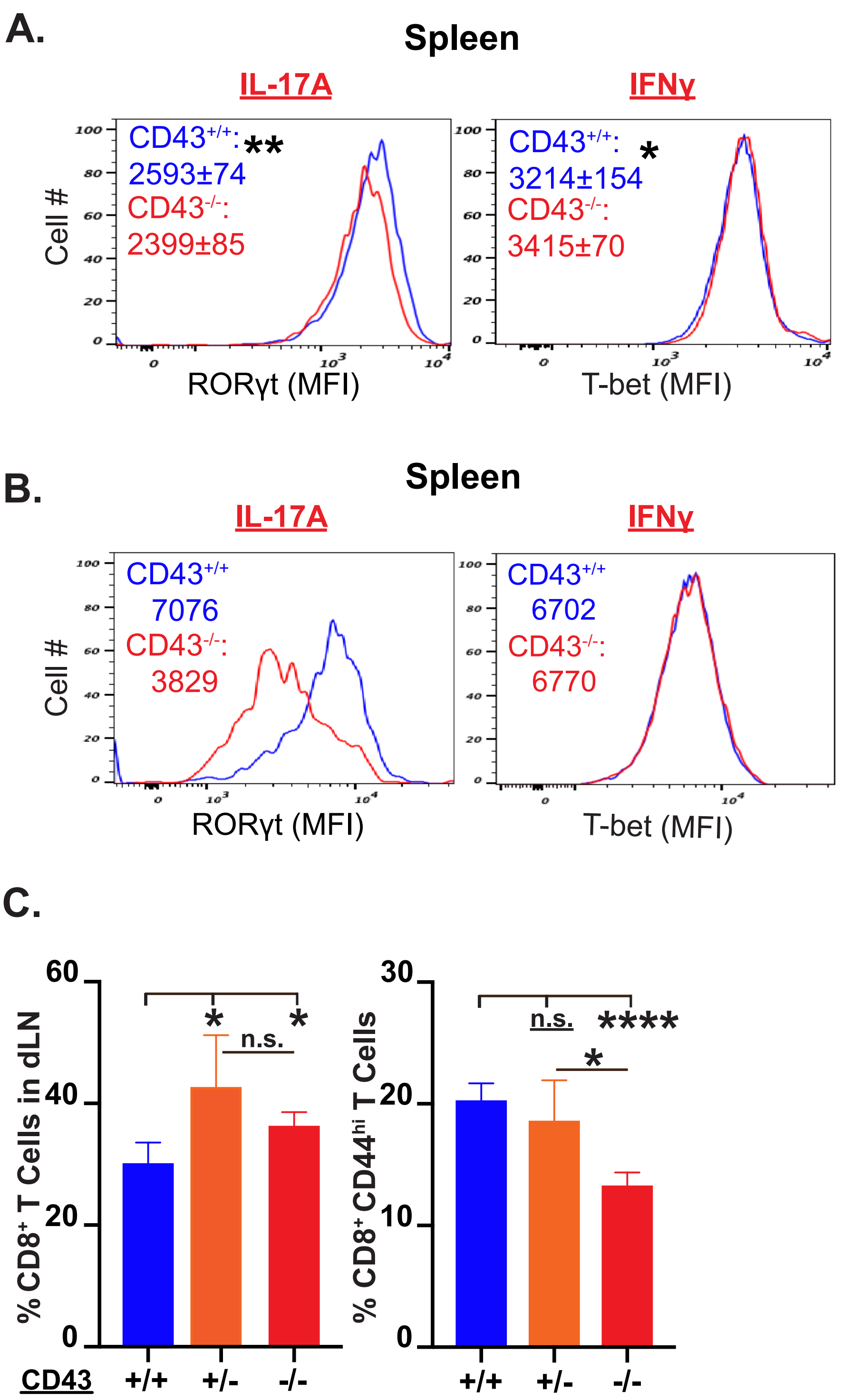

### Support Figure 4

#### Supplementary Figure 4
#### related to Figure 3.

## A.

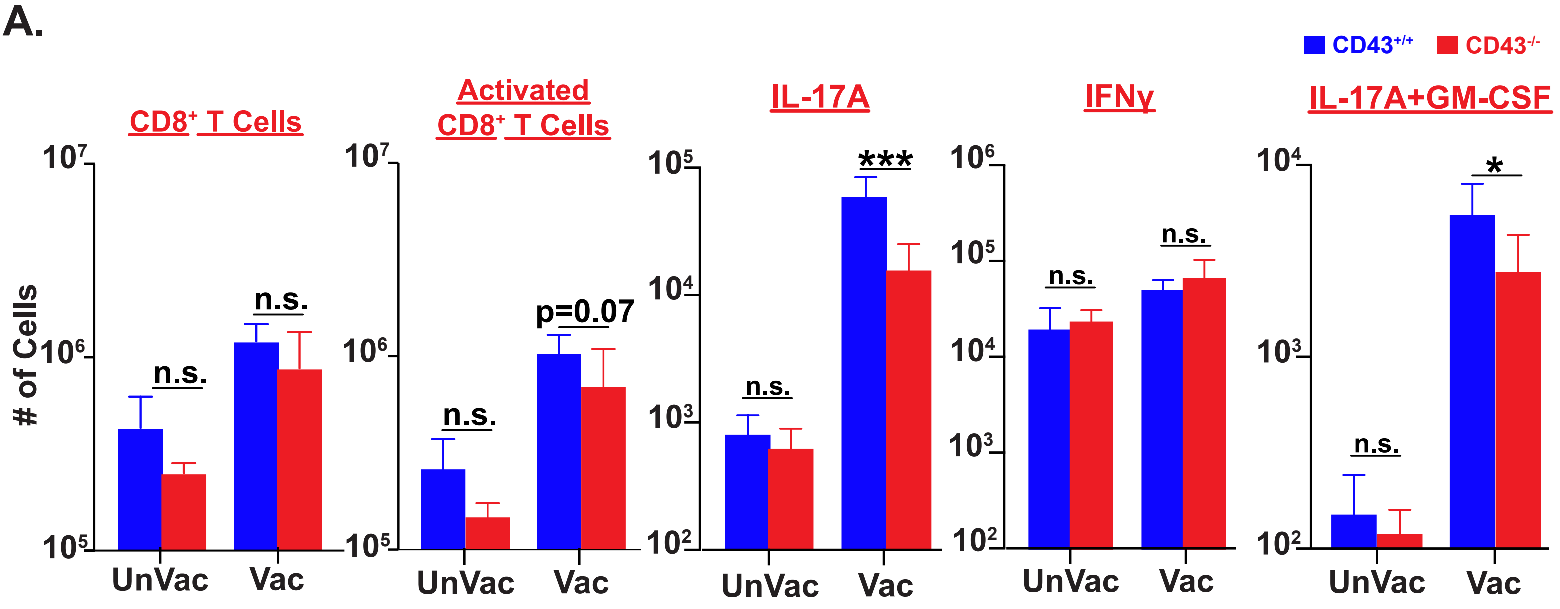


##### B. related to Discussion

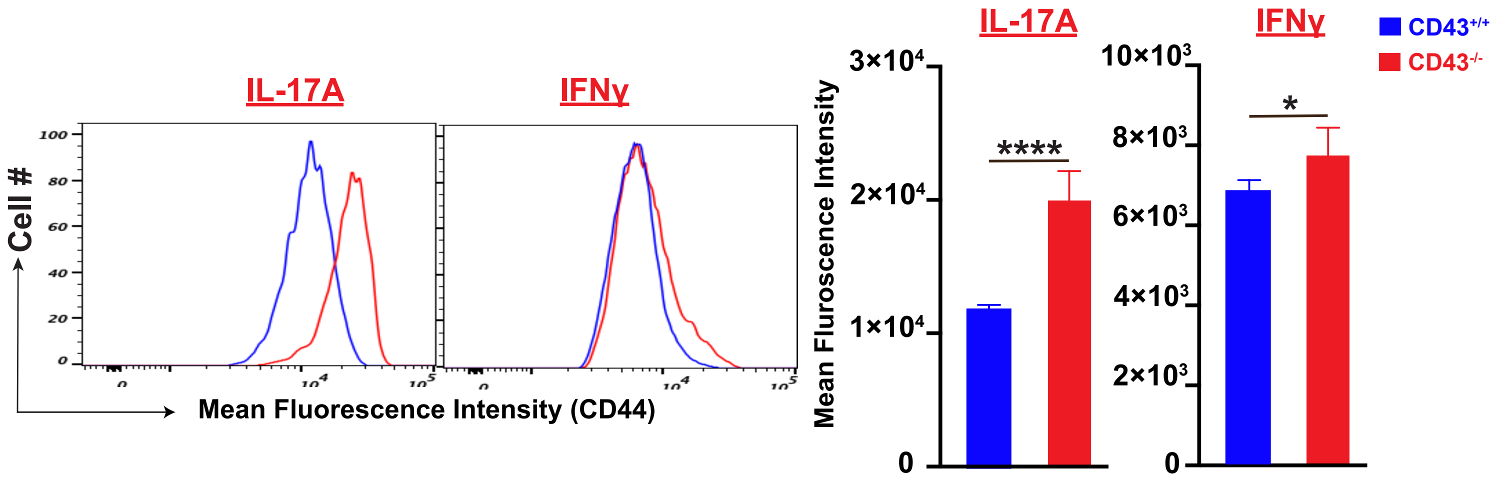
