## Supplemental Figure Legends for "Sialophorin is an essential host element for vaccine immunity against pulmonary fungal infections"

### SUPPLEMENTAL INFORMATION

**Supplementary Figure 1: CD43 preferentially expressed on fungal antigen-specific $CD8^+$ T cells**

(A) Glycosylated CD43 is preferentially expressed on Tc17 cells. Naïve 6-8-wk-old $CD43^{+/+}$ mice vaccinated with *Blastomyces dermatitidis #55*, and at indicated times (~4-wk and ~10-wk) single-cell suspensions from dLN and spleen were restimulated with anti-mouse CD3e and CD28 mAbs for 5hr in the presence of GolgiStop (BD Bioscience). Cells were surface stained before staining for intracellular cytokines and analyzed by flow cytometry. Data is representative of 1-2 independent experiments. Values are mean ± SD. N=3-6 mice/time. p**≤0.01 and p****≤0.0001 of $CD8^+$ T cells producing different cytokines.

(B) Glycosylated CD43 is preferentially expressed on fungal antigen-specific $CD8^+$ T cells. The single-cell suspensions (A) were used to enrich $CD8^+$ T cells (BD Bioscience Enrichment Kit) and co-incubated with heat-killed yeast-pulsed BMDC for 16hr with the last 5hr with GolgiStop (BD Bioscience). Cells were washed, surface and intracellular stained, and $CD43^+$ $CD8^+$ T cell responses were analyzed by flow cytometry. Data is representative of 1-2 independent experiments. Values are mean ± SD. N=3-6 mice/time. p*≤0.05, p**≤0.01, p***≤0.001, and p****≤0.0001 of $CD8^+$ T cells producing different cytokines.

**Supplementary Figure 2: CD43 for vaccine induced $CD8^+$ T cell responses**

(A-E) Naïve mice were vaccinated (#55, ~1x$10^5$ CFUs, by s.c.) and at ~3wks post-vaccination, single-cell suspensions were restimulated with anti-mouse CD3e and CD28 mAbs for 5hr in the presence of GolgiStop (BD Bioscience). $CD8^+$ T cells and their activation phenotype ($CD44^{hi}$)/cytokine expression in the spleen were measured by flow cytometry **(A-C).** Absolute numbers of cytokine-producing $CD8^+$ T cells in spleen and dLNs **(D-E)**. Data is representative of

5 independent experiments. Values are mean ± SD of N=4-5 mice/ group. $p^{*} \leq 0.05$, $p^{**} \leq 0.01$, and $p^{***} \leq 0.001$ of T cells in WT and KO mice.

**Supplementary Figure 3: CD43 for differentiation of effector $CD8^{+}$ T cells**

(A) Naïve $CD43^{+/+}$ and $CD43^{-/-}$ mice were vaccinated with #55 (~$1x10^{5}$ CFUs, s.c.) and at ~3wks post-vaccination, single-cell suspensions from spleen were restimulated with anti-mouse CD3e and CD28 mAbs for 5hr in the presence of GolgiStop (BD Bioscience). Cells were surface stained before staining for intracellular cytokines and transcription factors using eBioscience Transcription Factors Kit (Invitrogen). Transcription factor expression in cytokine-producing $CD8^{+}$ T cells were analyzed by flow cytometry. Data is representative of 2 independent experiments. Values are Mean Fluorescence Intensities ± SD of 4-5 mice/group. $p^{*} \leq 0.05$ and $p^{**} \leq 0.01$of T cells of WT versus KO mice.

(B) Enriched naïve $CD43^{+/+}$ and $CD43^{-/-}$ $CD8^{+}$ T cells were *co-transferred* (1:1) into naïve $TCR\alpha^{-/-}$ recipients. A day later, recipients were vaccinated and at ~3wks, single-cell suspensions of splenocytes were restimulated with anti-mouse CD3e and CD28 mAbs for 5hr in the presence of GolgiStop (BD Bioscience). Cells were surface stained before staining for intracellular cytokines and transcription factors using eBioscience Transcription Factors Kit (Invitrogen). Transcription factor expression in cytokine-producing $CD8^{+}$ T cells from *pooled samples* were analyzed by flow cytometry. Data is representative of 2 independent experiments. Values are Mean Fluorescence Intensities.

(C) Naïve $CD43^{+/+}$, $CD43^{+/-}$, and $CD43^{-/-}$ mice were vaccinated with #55 (~$1x10^{5}$ CFUs, s.c.) and at ~3wks post-vaccination, single-cell suspensions from dLN were restimulated with anti-mouse CD3e and CD28 mAbs for 5hr in the presence of GolgiStop (BD Bioscience). Cells were surface stained and activated/$CD8^{+}$ T cells were analyzed by flow cytometry. Data is representative of 2 independent experiments. Values are mean ± SD of 5 mice/group. p*≤0.05 and p****≤0.0001of T cells of WT vs Hets vs KO mice.

**Supplementary Figure 4: CD43 for recall responses and expression of CD44**

(A) Sialophorin is required for vaccine-mediated immunity. Naïve $CD43^{+/+}$ and $CD43^{-/-}$ mice were vaccinated, rested for ~6 weeks, and intratracheally challenged with lethal yeast #26199 along with unvaccinated controls. Single-cell suspensions from lungs of cohorts of vaccinated and challenged mice were restimulated with anti-mouse CD3e and CD28 mAbs for 5hr and $CD8^+$ T cell responses were analyzed by flow cytometry. Data is representative of two independent experiments. Values are mean ± SD. N=5 mice/group. p*≤0.05 and p***≤0.001 of T cells of unvaccinated and vaccinated WT/KO mice.

(B) Sialophorin deficiency leads to high CD44 expression. Splenocytes from the vaccinated mice were harvested at 3 weeks post-vaccination and CD44 expression on cytokine-producing $CD8^+$ T cells was analyzed by flow cytometry. Data is representative of five independent experiments. Values are mean ± SD. N=5 mice/group. p*≤0.05 and p***≤0.001 of WT/KO mice.
